# A novel deep proteomic approach in human skeletal muscle unveils distinct molecular signatures affected by aging and resistance training

**DOI:** 10.1101/2023.06.02.543459

**Authors:** Michael D. Roberts, Bradley A. Ruple, Joshua S. Godwin, Mason C. McIntosh, Shao-Yung Chen, Nicholas J. Kontos, Anthony Agyin-Birikorang, J. Max Michel, Daniel L. Plotkin, Madison L. Mattingly, C. Brooks Mobley, Tim N. Ziegenfuss, Andrew D. Fruge, Andreas N. Kavazis

**Affiliations:** School of Kinesiology, Auburn University, Auburn, AL USA; Seer, Inc., Redwood City, CA, USA; The Center for Applied Health Sciences, Canfield, OH, USA; College of Nursing, Auburn University, Auburn, AL, USA

**Author notes:** Address correspondence to: Michael D. Roberts, PhD Professor, School of Kinesiology Director, Nutrabolt Molecular and Applied Sciences Laboratory.

**Keywords:** skeletal muscle, deep proteomics, aging, resistance training

## Abstract

We examined the myofibrillar (MyoF) and non-myofibrillar (non-MyoF) proteomic profiles of the vastus lateralis (VL) muscle of younger (Y, 22±2 years old; n=5) and middle-aged participants (MA, 56±8 years old; n=6), and MA following eight weeks of knee extensor resistance training (RT, 2d/week). Shotgun/bottom-up proteomics in skeletal muscle typically yields wide protein abundance ranges that mask lowly expressed proteins. Thus, we adopted a novel approach whereby the MyoF and non-MyoF fractions were separately subjected to protein corona nanoparticle complex formation prior to digestion and Liquid Chromatography Mass Spectrometry (LC-MS) analysis. A total of 10,866 proteins (4,421 MyoF and 6,445 non-MyoF) were identified. Across all participants, the number of non-MyoF proteins detected averaged to be 5,645±266 (range: 4,888–5,987) and the number of MyoF proteins detected averaged to be 2,611±326 (range: 1,944–3,101). Differences in the non-MyoF (8.4%) and MyoF (2.5%) proteome were evident between age cohorts. Further, most of these age-related non-MyoF proteins (447/543) were more enriched in MA versus Y. Several biological processes in the non-MyoF fraction were predicted to be operative in MA versus Y including (but not limited to) increased cellular stress, mRNA splicing, translation elongation, and ubiquitin-mediated proteolysis. Non-MyoF proteins associated with splicing and proteostasis were further interrogated, and in agreement with bioinformatics, alternative protein variants, spliceosome-associated proteins (snRNPs), and proteolysis-related targets were more abundant in MA versus Y. RT in MA non-significantly increased VL muscle cross-sectional area (+6.5%, p=0.066) and significantly increased knee extensor strength (+8.7%, p=0.048). However, RT modestly altered the MyoF (∼0.3%, 11 upregulated and two downregulated proteins) and non-MyoF proteomes (∼1.0%, 56 upregulated and eight downregulated proteins, p<0.01). Further, RT did not affect predicted biological processes in either fraction. Although participant numbers were limited, these preliminary results using a novel deep proteomic approach in skeletal muscle suggest that aging and RT predominantly affects protein abundances in the non-contractile protein pool. However, the marginal proteome adaptations occurring with RT suggest either: a) this may be an aging-associated phenomenon, b) more rigorous RT may stimulate more robust effects, or c) RT, regardless of age, subtly affects skeletal muscle protein abundances in the basal state.

## INTRODUCTION

Aging adversely affects skeletal muscle physiology as evidenced by a reduction in muscle stem (or satellite) cell content, a loss of myofibrillar protein, and a loss in motor units and myofibers (1–4). Muscle aging is also associated with impairments in mitochondrial function, dysfunctional redox balance, and altered proteostasis (5–7). The culmination of these events likely contributes to a loss in muscle mass, which according to a recent review, is accelerated in all the body’s musculature past the age of 50 years old (8).

Resistance training can reverse certain aspects of skeletal muscle aging. For instance, weeks to months of resistance training in older participants has been shown to increase tissue-level and myofiber hypertrophy and muscle satellite cell content (9). Resistance training also catalyzes skeletal muscle mitochondrial biogenesis and remodeling in older participants (10–14), and weeks to months of resistance training alters nuclear and mitochondrial DNA methylation patterns in older participants which may lead to “rejuvenating” effects on global mRNA expression patterns (11, 15, 16).

Proteomic investigations intend to examine the entire detectable protein expression signature of a given tissue under various experimental conditions. While other -omics-based approaches exist (i.e., genomics, epigenomics, transcriptomics, and metabolomics), it has been posited that proteomic signatures likely best translate to cellular and tissue phenotypes (17). Past proteomic investigations have provided tremendous insight as to how myofiber type, aging, and exercise training affect the skeletal muscle molecular milieu (1, 14, 18–22). Notwithstanding, skeletal muscle-based proteomics poses technical challenges. For example, skeletal muscle tissue processing with general lysis buffers results in the clearance of insoluble (e.g., contractile) proteins (23), and if standard bottom-up proteomics is employed thereafter, the relative abundances of these proteins will ultimately be inaccurate. Even if care is taken in isolating the poorly soluble contractile and soluble non-contractile protein fractions, another pitfall lies in lowly-abundant proteins being masked by highly abundant proteins in each fraction (1). Single fiber isolation techniques have increased the depth of proteins detected (21, 24). However, certain disadvantages with this method exist including the burdensome process of tissue digestion and fiber dissection, the need for pooling myofibers to obtain adequate protein for proteomics, and the inability to detect proteins enriched in the extracellular matrix or stromal cells.

A novel deep proteomics approach in human plasma was recently published whereby unique nano-bio interaction properties of multiple magnetic nanoparticles (NPs) was leveraged for automated protein separation (referred to as the Proteograph assay; Seer, Inc. Redwood, CA, USA) (25). Downstream digestion followed by liquid chromatograph coupled to mass spectrometry (LC-MS) analyses enabled the identification of over 2,000 plasma proteins and this provided approximately a 10-fold increase in depth compared to prior studies that utilized other strategies to deplete plasma of highly abundant proteins (26, 27). However, this approach has not been performed in human skeletal muscle. Thus, we sought to leverage this technology, along with our prior method of muscle tissue fractionation (23), to examine the proteomic signatures of the myofibrillar (MyoF) and non-myofibrillar (non-MyoF) fractions from the vastus lateralis (VL) muscle of a subset of younger (Y, 22±2 years old, n=5) and middle-aged (MA, 56±8 years old, n=6) participants. We also sought to determine how eight weeks of unilateral knee extensor resistance training affected the MyoF and non-MyoF proteomic signatures in the MA cohort. Given some of our past work in this area (1), we hypothesized that more non-MyoF proteins would be altered by aging when comparing MA and Y participants. We also hypothesized that resistance training in MA participants would affect more non-MyoF versus MyoF proteins. However, we did not adopt an *a priori* hypothesis regarding which proteins or biological processes would be affected between comparisons given the novelty of interrogating skeletal muscle using the Proteograph assay.

## METHODS

### Ethical approval and study design

Muscle specimens were obtained from two studies whereby approval was obtained from the Auburn University Institutional Review Board. The first protocol in untrained MA participants (approved protocol #21-461 MR 2110) involved investigating the effects of a dietary supplement (312 mg of combined Wasabia japonica extract, theacrine, and copper (I) niacin chelate) versus a placebo on potential blood marker responses over an eight-week period. A unilateral leg resistance training (two days/week) protocol was implemented to perform non-supplementation secondary analyses as presented herein. The six MA participants included in the current study were in the placebo group; thus, no confounding effects of dietary supplementation were expected. Y participant muscle tissue was banked from a prior study examining how ten weeks of daily peanut protein supplementation affected resistance training outcomes in untrained individuals (approved protocol #19–249 MR 1907) (28). Notably, muscle tissue from these participants was collected in the basal state prior to the intervention. Hence, again, there were no potential confounding effects of supplementation. Study procedures for both projects were in accordance with the most recent revisions of the Declaration of Helsinki except for the MA study not being pre-registered as a clinical trial.

### Study Design and Training Paradigm in MA participants

#### Knee extensor resistance training

The resistance training intervention consisted of supervised unilateral leg extensions (two days/week for eight weeks), and the intervention was preceded and followed by strength and VL muscle assessments (described in later paragraphs). All MA participants trained their right legs whereby each training session consisted of five sets of 12 repetitions. The beginning training load was established at ∼40% of the participants’ three-repetition maximum (3RM). After each set, participants verbally articulated their perceived repetitions in reserve (RIR) (29), and training load was adjusted accordingly. RIR values of 0-2 after a set resulted in no training load change in each session. RIR values of 3-5 for consecutive sets resulted in the training load being increased by 5-10%. For RIR values ≥6 after one set, the training load was increased by 10-20%. If the weight could not be performed with full range of motion, or the participant could not complete 12 repetitions for a given set, the training load was decreased accordingly.

#### Strength testing

The first and last workout of the eight-week training paradigm consisted of maximal leg extensor-flexion torque assessments using isokinetic dynamometry (Biodex System 4; Biodex Medical Systems, Inc., Shirley, NY, USA) and 3RM leg extensor strength testing. Prior to dynamometer testing, the participant’s lateral epicondyle was aligned with the axis of the dynamometer’s lever arm, and the hip was positioned at 90°. The participant’s shoulders, hips, and leg were strapped and secured for isolation during testing. Following three warm-up trials at a submaximal effort, participants completed five maximal voluntary isokinetic knee extension and flexion actions at 60 degrees/second. Participants were provided verbal encouragement during each contraction. The isokinetic contraction resulting in the greatest peak torque value was used for analyses. Approximately five minutes following isokinetic dynamometry testing, participants performed 3RM strength testing using a free-weight apparatus. Prior to testing, participants were given a warm-up load and instructed to complete 10 repetitions. After participants recorded their RIR for the warmup set, the weight was adjusted accordingly for another warm-up set of five repetitions. RIR was recorded again to determine the participants starting load for a 3RM attempt. The load was incrementally increased 5-10% per 3RM attempt until 3RM testing concluded, indicated by failure of full range of motion on any of the repetitions, or if RIR recorded was 0. Participants were allowed a full three minutes of recovery between attempts. The isokinetic dynamometry and 3RM testing described was similar for both the first and final workout.

### Testing Sessions in MA participants

#### Urine specific gravity testing for hydration

Participants performed a testing battery prior to the start of training (PRE) and 3-5 days following the last resistance training workout (POST). Participants arrived for testing at a minimum of 4 hours fasted and well hydrated. Upon arrival participants submitted a urine sample (∼5 mL) for urine specific gravity assessment (USG). Measurements were performed using a handheld refractometer (ATAGO; Bellevue, WA, USA). USG levels in all participants were ≤ 1.020, indicating sufficient hydration (30).

#### Body composition testing

Body composition was assessed using multi-frequency bioelectrical impedance analysis (InBody 520, Biospace, Inc. Seoul, Korea). From the scan, body fat percentage was recorded. Previously determined test-retest reliability yielded an intraclass correlation coefficient (ICC_3,1_) of 0.99, standard error of the measurement (SEM) of 0.87%, and minimal difference (MD) of 1.71% for body fat percentage.

#### Ultrasonography assessment for muscle morphology

A detailed description of VL assessments using ultrasonography has been published previously by our laboratory (31, 32). Briefly, real-time B-mode ultrasonography (NextGen LOGIQe R8, GE Healthcare; Chicago, IL, USA) using a multifrequency linear-array transducer (L4-12T, 4–12 MHz, GE Healthcare) was used to capture VL muscle cross-sectional area (mCSA). Prior to scans, the mid-thigh location was determined by measuring the total distance from the mid-inguinal crease in a straight line to the proximal patella, with the knee and hip flexed at 90°, a mark was made using a permanent marker at 50% of the total length. From that location, a permanent marker was used transversely to mark the mid-belly of the VL. This marking is where all pre-intervention ultrasound images were taken as well as the muscle biopsy (described below). All post-intervention images were taken at the pre-intervention biopsy scar to ensure location consistency between scans. During mCSA scans, a flexible, semirigid pad was placed around the thigh and secured with an adjustable strap to allow the probe to move in the transverse plane. Using the panoramic function of the device (LogicView, GE Healthcare), images were captured starting at the lateral aspect of the VL and moving medially until rectus femoris was visualized, crossing the marked location. All ultrasound settings were held constant across participants and laboratory visits (frequency: 10 MHz, gain: 50 dB, dynamic range: 75), and scan depth was noted and held constant across time points per participant. Images were downloaded and analyzed offline using ImageJ software (National Institutes of Health, Bethesda, MD, USA). All ultrasound images were captured and analyzed by the same investigators at each timepoint. Previously determined test-retest reliability on 10 participants measured twice within 24 hours (where BAR captured images and JSG analyzed images) yielded an intraclass correlation of 0.99 and standard error of measurement of 0.60 cm^2^.

#### Collection of muscle tissue

Muscle biopsies from all participants were obtained from the mid-belly of the right VL, and sampling time of day was standardized for MA participants at pre and post resistance training intervention. Lidocaine (1%, 1.0 mL) was injected subcutaneously above the skeletal muscle fascia at the previously marked location. After five minutes of allowing the anesthetic to take effect, a small pilot incision was made using a sterile Surgical Blade No. 11 (AD Surgical; Sunnyvale, CA, USA), and the 5-gauge biopsy needle was inserted into the pilot incision ∼1 cm below the fascia. Approximately 30-50 mg of skeletal muscle was removed using a double chop method and applied suction. Following biopsies, tissue was rapidly teased of blood and connective tissue, placed in pre-labeled foils, flash frozen in liquid nitrogen, and subsequently stored at −80°C until processing described below.

### MyoF and non-MyoF protein fractionation

The MyoF and non-MyoF protein fractions were isolated per methods published by our laboratory and others (23, 33). On the day of homogenization, muscle tissue was powdered on a liquid nitrogen-cooled ceramic mortar and pestle. Approximately 30 mg of tissue was homogenized using tight-fitting pestles in 500 µL of 25 mM Tris, pH 7.2, 0.5% Triton X-100, with added protease inhibitors (Promega, cat# G6521; Madison, WI, USA). Samples were centrifuged at 1,500 g for 10 minutes at 4°C, supernatants (non-MyoF fraction) were transferred to new 1.7 mL tubes, and tubes were stored at −80°C until shipment on dry ice to Seer, Inc. Remaining MyoF pellets were kept on ice and thoroughly aspirated with micro-pipet tips to remove residual supernatant. Thereafter, 300 μL of solubilization buffer was added which contained 20 mM Tris-HCl, pH 7.2, 100 mM KCl, 20% glycerol, 1 mM DTT, 50 mM spermidine with added protease inhibitors (Promega, cat# G6521). Samples were then homogenized using tight-fitting pestles and stored at −80°C until shipment on dry ice to Seer, Inc.

### MyoF and non-MyoF Proteograph assays and proteomics

#### Proteograph assay

Proteomics analysis was performed at Seer, Inc. (Redwood City, CA, USA). For each sample, 250 μL of received sample was subjected to the Seer Proteograph Assay protocol. After loading samples onto the SP100 Automation Instrument, protein corona formation and processing was initiated to generate desalted purified peptides for protein identification using Reversed Phase (RP) LC-MS. To form the protein corona, Seer’s proprietary NPs were mixed with the samples and incubated at 37°C for 1 hour. Unbound proteins were removed prior to downstream wash, reduction, alkylation, and protein digestion steps which were performed according to Seer’s Proteograph Assay protocol (25).

#### LC-MS configuration

Peptides obtained from each of the five NP mixtures were separately reconstituted according in a solution of 0.1% formic acid and 3% acetonitrile (34) spiked with 5 fmol μL PepCalMix from SCIEX (Framingham, MA, USA). Reconstitution volumes varied by NP types to allow for constant peptide quantity for MS injection between samples regardless of starting volume (240 ng: NP1, 400 ng: NP2, 360 ng: NP3, 120 ng: NP4, and 320 ng: NP5). 4 µL of each sample were analyzed with a Ultimate3000 RLSCnano LC system coupled with a Orbitrap Fusion Lumos mass spectrometer (Thermo Fisher; Waltham, MA, USA). Peptides were loaded on an Acclaim PepMap 100 C18 (0.3 mm ID × 5 mm) trap column and then separated on a 50 cm μPAC analytical column (PharmaFluidics, Belgium) at a flow rate of 1 μL/min using a gradient of 5–25% solvent B (100% ACN) mixed into solvent A (100% water) over 26 minutes. The mass spectrometer was operated in Data Independent Acquisition (DIA) mode using 10 m/z isolation windows from 380-1200 m/z and 3-second cycle time. MS1 scans were acquired at 60k resolution and MS2 at 30k resolution.

#### Data Processing

DIA LC-MS data were processed using Proteograph Analysis Suite (PAS) v2.1 (Seer, Inc) using the DIA-NN search engine (version 1.8.1) in library-free mode searching MS/MS spectra against an in silico predicted library based on Uniprot’s Homo Sapiens reference database (UP000005640_9606, download December 9, 2022). Library-free search parameters included trypsin digestion allowing for one missed cleavage, N-terminal methionine excision, fixed modification of cysteine carbamidomethylation, peptide length of 7-30 amino acids, precursor range of 300-1800 m/z, and fragment ion range of 200-1800 m/z. Heuristic protein inference was enabled, MS1 and MS2 mass accuracy was set to 10 ppm. Precursor FDR was set to 0.01, and PG q-value was set to 0.01. Quantification was performed on summed abundances of all unique peptides considering only precursors passing the q-value cutoff. PAS summarizes all NP values for a single protein into a single quantitative value. Specifically, a single protein may have been measured up to five times, once for each nanoparticle. To derive the single measurement value, PAS uses a maximum representation approach, whereby the single quantification value for a particular peptide or protein group represents the quantitation value of the NP which most frequently has measured any given proteins across all samples.

The relative abundances of protein targets were obtained by normalizing raw spectra values for each identified protein to total spectra within-subject. After normalization, undetected protein abundance values were set at zero. Protein values are presented as spectra-normalized values in all figures and results.

### Statistics and bioinformatics

Data processing and statistical analysis was performed using Microsoft Excel for Microsoft 365 (Redmond, WA, USA) and GraphPad Prism version 9.2.0 (San Diego, CA, USA). Independent samples t-tests were used for Y versus MA (pre-intervention) to determine age effects, and dependent samples t-tests were used to determine training effects in MA. All data in tables and figures are presented as mean ± standard deviation (SD) values. Training phenotypes were considered significantly different at p<0.05, although approaching values (i.e., p<0.100) were discussed as “numerical” changes due to limited n-sizes. Conversely, significant aging and training effects for protein targets were established as p<0.01 for enhanced stringency given the high number of identified proteins, although again approaching values (i.e., p<0.05) were discussed in certain circumstances due to limited n-sizes.

Bioinformatics was performed using PANTHER v17.0 (35, 36). First, protein lists from each fraction were characterized using the functional classification tool. Next, overrepresentation tests of PANTHER GO-Slim biological processes were performed between Y and MA participants and in MA participants from pre-to-post training. Parameters for statistical overrepresentation tests included the following: i) entered proteins had to meet the aforementioned p<0.01 significance threshold, ii) protein lists were entered separately based on being up- or downregulated to generate a list of biological processes that were predicted to be directionally affected, and iii) Fisher tests with Bonferroni adjusted p<0.05 values were used as significance thresholds.

## RESULTS

### MA versus Y phenotypes, and MA responses to eight weeks of resistance training

MA were significantly older than Y participants (Y: 22 ± 2 years old, MA: 56 ± 8 years old; p<0.001). However, compared to Y participants, pre-intervention MA participant body mass (Y: 69.9 ± 14.3 kg, MA: 76.6 ± 14.7 kg; p=0.374), percent body fat (Y: 35.5 ± 4.6 %, MA: 27.6 ± 6.6 %; p=0.071), and VL mCSA (Y: 19.0 ± 2.7 cm^2^, MA: 18.4 ± 5.2 cm^2^; p=0.818) were not significantly different.

In MA participants, the eight-week training protocol non-significantly increased VL mCSA (PRE: 18.4 ± 5.2 cm^2^, POST: 19.6 ± 4.2 cm^2^; p=0.066) and significantly increased isokinetic knee extensor strength at 60 degrees/s (PRE: 141 ± 79 N•m, POST: 153 ± 71 N•m; p=0.048).

### Characteristics of proteins identified in the MyoF and non-MyoF fractions

A total of 6,445 non-MyoF proteins and 4,421 MyoF proteins were identified in at least one participant (Fig. 1a). Across all participants, the number of non-MyoF proteins detected averaged to be 5,645 ± 266 (range: 4,888–5,987) and the number of MyoF proteins detected averaged to be 2,611 ± 326 (range: 1,944–3,101).

**Figure 1.**
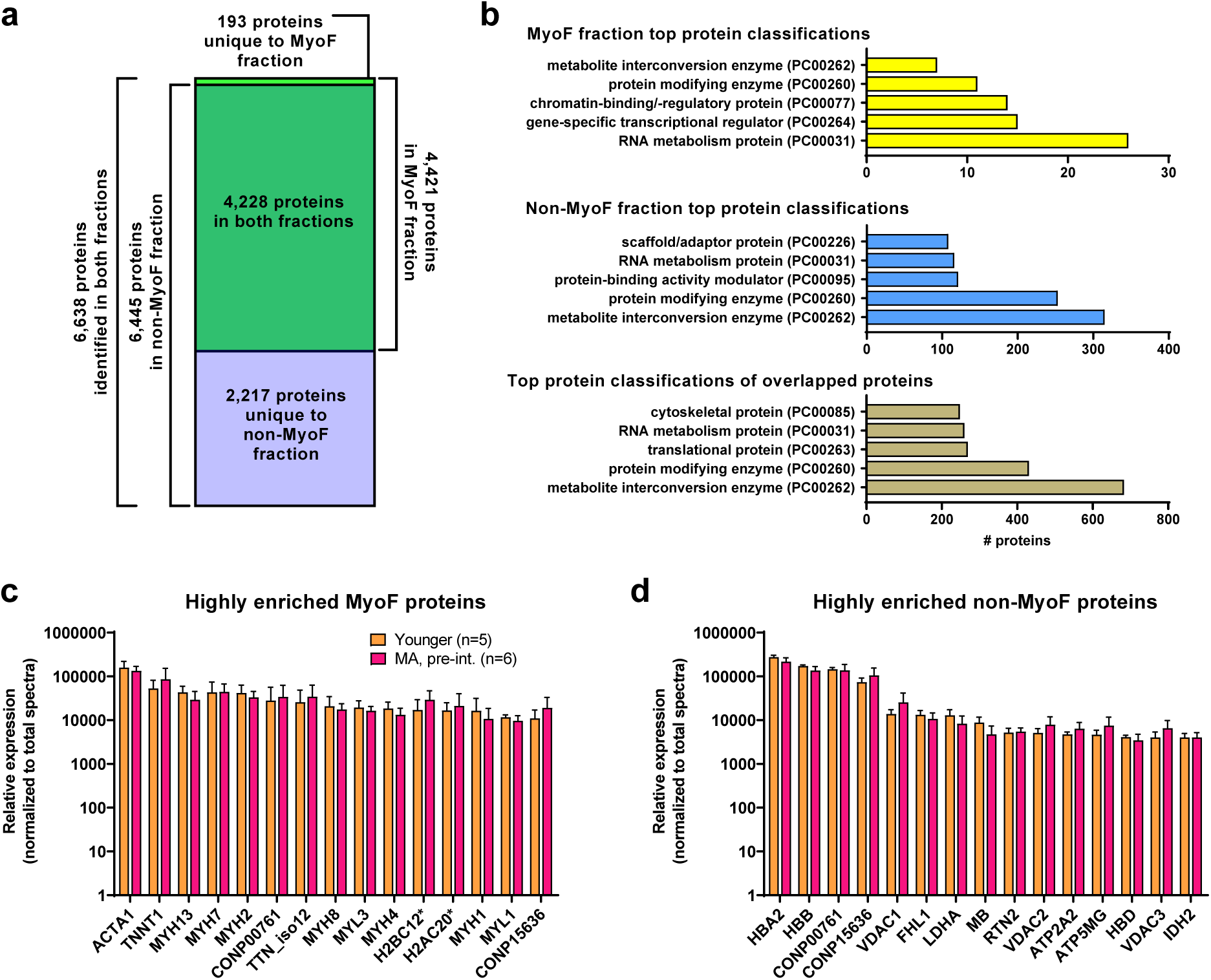
MyoF and non-MyoF protein characteristics. Legend: Data presented for MA (pre-intervention) and Y include the total number of proteins identified in each fraction (panel a), the top 5 protein classifications from each fraction (panel b), the top 15 highly enriched MyoF proteins (panel c), and the top 15 highly enriched non-MyoF proteins (panel d). Data in panels c and d are presented as means with standard deviation bars, and y-axes were scaled as log_10_ for improved visualization. Symbols: *, indicates multiple histone isoforms were congregated into these two targets based on sequence similarities. Protein names for gene symbols in panel c: ACTA1, Actin Alpha 1, Skeletal Muscle; TNNT1, Troponin T1; MYH13/7/2/8/4/1, myosin heavy chain isoforms 13/7/2/8/4/1; TTN_iso12, titin, isoform 12; MYL3/1/, myosin light chain isoforms 3/1; H2BC12, Histone H2B type 1-K/C/E/F/G/I/type F-S; H2AC20, Histone H2A type 2-A/C. Protein names for gene symbols in panel d: HBA2, Hemoglobin subunit alpha; HBB, Hemoglobin subunit beta; VDAC1, Voltage-dependent anion-selective channel protein 1; FHL1, Four and a half LIM domains protein 1; LDHA, L-lactate dehydrogenase A chain; MB, myoglobin; RTN2, Isoform RTN2-C of Reticulon-2; VDAC2, Voltage-dependent anion-selective channel protein 2; ATP2A2, Sarcoplasmic/endoplasmic reticulum calcium ATPase 2; ATP5MG, ATP synthase subunit g, mitochondrial; HBD, Hemoglobin subunit delta; VDAC3, Voltage-dependent anion-selective channel protein 3; IDH2, Isocitrate dehydrogenase [NADP], mitochondrial. Other note: CONP00761/15636 are non-annotated proteins found in both fractions.

A total of 4,228 proteins overlapped in both fractions yielding 2,217 unique non-MyoF proteins, 193 unique MyoF proteins, and 6,638 unique proteins identified. Using the PANTHER Classification System classifications, the top five protein classes of MyoF proteins, non-MyoF proteins, and proteins in both fractions are presented in Fig 1b. The top 15 enriched MyoF and non-MyoF proteins in MA (pre-intervention) and Y are presented in Figure 1 (panels c and d). None of the 15 MyoF or non-MyoF proteins met the p<0.01 significance criteria between age cohorts.

### MYH isoform peptide identification information

Myosin heavy chain isoforms have been intensely studied in human skeletal muscle for fiber typing purposes and prominent isoforms include the slow-twitch type I isoform (encoded by the MYH7 gene) as well as the fast-twitch IIA (encoded by the MYH2 gene) and IIX (encoded by the MYH1 gene) isoforms (37). However, other MYH isoforms were highly enriched in the MyoF fraction according to data presented in Fig. 1c. Because of this, we opted to provide the peptide sequences used for detecting some of these isoforms in Table 1 below.

**Table 1.** Peptide sequences of highly enriched myosin heavy chain isoforms in the MyoF fraction. Legend: these data contain the peptide sequences used for alignment to identify the several highly abundant myosin heavy chain isoforms in the MyoF fraction.

| Myosin heavy chain protein (gene) | Uniprot ID | Peptide sequence<br>(location; total length of protein) |
| --- | --- | --- |
| Myosin heavy chain I (MYH7) | P12883 | TKYETDAIQR<br>(amino acids 1373-1382; 1935) |
| Myosin heavy chain IIa (MYH2) | Q9UKX2 | TLAQLFSGAQTAEGEGAGGGAK<br>(amino acids 619-640; 1941) |
| Myosin heavy chain-perinatal (MYH8) | P13535 | LAQIITR<br>(amino acids 784-790; 1937) |
| Myosin IIb (MYH4) | Q9Y623 | TLEDQLSEIK<br>(amino acids 1255-1264; 1939) |
| Myosin heavy chain IIx (MYH1) | P12882 | TEAGATVTVK<br>(amino acids 64-73; 1939) |

### Additional MyoF and non-MyoF protein characteristics

Aside from using PANTHER protein classifications, we wanted to present the depth of proteins detected based on processes relevant to muscle biology. Through manual interrogation we found that both fractions contained proteins found in nuclei that regulate chromatin structure and gene expression (Fig. 2a), mitochondrial proteins related to oxidative phosphorylation and mitochondrial protein synthesis (Fig. 2b), proteolysis-related proteins (Fig. 2c), proteins localized to the sarcolemma and solute carrier proteins (Fig. 2d), exercise-relevant phosphosignaling proteins (Fig. 2e), and proteins associated with protein synthesis (Fig. 2f).

**Figure 2.**
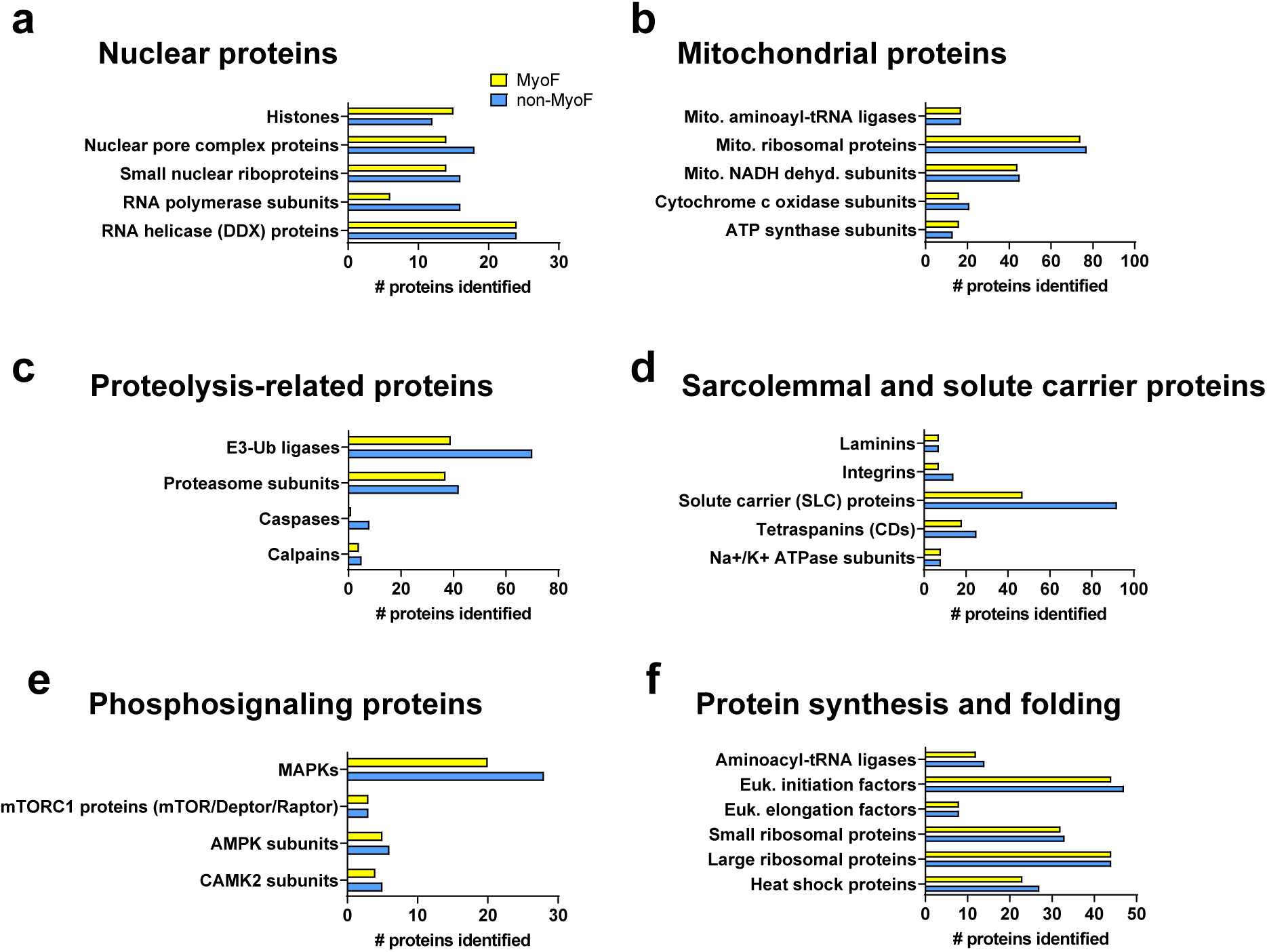
Additional protein classifications. Legend: Each fraction was found to contain nuclear proteins that regulate chromatin structure and gene expression (panel a), mitochondrial proteins related to oxidative phosphorylation and mitochondrial protein synthesis (panel b), proteolysis-related proteins (panel c), proteins localized to the sarcolemma and solute carrier proteins (panel d), exercise-relevant phosphosignaling proteins (panel e), and proteins associated with protein synthesis (panel f). Abbreviations: Mito., mitochondrial; Ub, ubiquitin; MAPKs, mitogen of activated protein kinases; mTORC1, mechanistic target of rapamycin complex 1; AMPK, AMP-activated protein kinase; CAMK, Ca^2+^/calmodulin-dependent protein kinase; Euk., eukaryotic.

### MyoF and non-MyoF protein differences in MA (pre-intervention) versus Y

In MA (pre-intervention) and Y participants, 112 of the 4,421 MyoF proteins met the p<0.01 significance threshold equating to ∼2.5% of the MyoF proteome being affected with aging. 111 of these 112 MyoF proteins were significantly greater in Y versus MA participants (see Table 2 for the top 15 proteins), and only one MyoF protein was significantly greater in the MA versus Y participants (TMPO, Lamina-associated polypeptide 2 isoforms beta/gamma).

**Table 2.** Top 15 of 111 MyoF proteins greater in Y versus MA participants. Legend: Data are presented as mean ± standard deviations for individual protein spectra values (normalized to total run spectra values) in younger (Y) and pre-intervention middle-aged (MA) participants. Protein targets are sorted from most to least abundant in the Y cohort.

| Protein (gene symbol) | Y (n=5) | MA (n=6) | p-value |
| --- | --- | --- | --- |
| Heat shock protein beta-1 (HSPB1) | 2062 ± 677 | 700 ± 380 | 0.0022 |
| Isoform 3 of ATP-dependent 6-phosphofructokinase (PFKM) | 700 ± 157 | 316 ± 169 | 0.0038 |
| Calmodulin-2 (CALM2) | 322 ± 94 | 146 ± 45 | 0.0026 |
| Protein NipSnap homolog 2 (NIPSNAP2) | 309 ± 62 | 201 ± 44 | 0.0079 |
| Protein-arginine deiminase type-2 (PADI2) | 306 ± 86 | 138 ± 25 | 0.0013 |
| Heat shock protein HSP 90-beta (HSP90AB1) | 254 ± 76 | 116 ± 62 | 0.0091 |
| Isoform 2 of Y-box-binding protein 3 (YBX3) | 203 ± 66 | 86 ± 33 | 0.0039 |
| Isoform 2 of 2,4-dienoyl-CoA reductase [(3E)-enoyl-CoA-producing], mitochondrial (DECR1) | 168 ± 51 | 64 ± 27 | 0.0020 |
| Isoform 2 of Phosphorylase b kinase regulatory subunit alpha (PHKA1) | 167 ± 51 | 73 ± 31 | 0.0044 |
| 28 kDa heat- and acid-stable phosphoprotein (PDAP1) | 146 ± 54 | 52 ± 38 | 0.0078 |
| Isoform 2 of Phosphorylase b kinase gamma catalytic chain (PHKG1) | 129 ± 43 | 52 ± 25 | 0.0048 |
| cAMP-dependent protein kinase catalytic subunit alpha (PRKACA) | 120 ± 44 | 46 ± 24 | 0.0063 |
| cAMP-dependent protein kinase type II-alpha regulatory subunit (PRKAR2A) | 118 ± 44 | 45 ± 23 | 0.0060 |
| NADH dehydrogenase [ubiquinone] iron-sulfur protein 8, mitochondrial (NDUFS8) | 115 ± 31 | 68 ± 15 | 0.0094 |
| Smoothelin-like protein 1 (SMTHL1) | 106 ± 43 | 43 ± 16 | 0.0087 |

Bioinformatics of the 111 MyoF proteins that were significantly greater in the Y versus MA participants indicated that no biological processes were predicted to be affected between age cohorts.

When performing the MA (pre-intervention) and Y participant comparisons for non-MyoF proteins, 543 proteins met the p<0.01 significance threshold equating to ∼8.4% of the non-MyoF proteome being affected with aging. 96 of these 543 non-MyoF proteins were significantly greater in Y versus MA participants, and 447 non-MyoF proteins were significantly greater in MA versus Y participants. Table 3 contains the top 15 proteins that showed differential abundances between Y versus MA participants.

**Table 3.** Top 30 of 543 non-MyoF proteins different between Y versus MA participants. Legend: Data are presented as mean ± standard deviations for individual protein spectra values (normalized to total run spectra values) in younger (Y) and pre-intervention middle-aged (MA) participants. Protein targets more abundant in Y versus MA are sorted from most to least abundant in the Y cohort. Protein targets more abundant in MA versus Y are sorted from most to least abundant in the MA cohort.

| Protein (gene symbol) | Y (n=5) | MA, pre- (n=6) | p-value |
| --- | --- | --- | --- |
| *Top 15 of 96 non-MyoF proteins higher in Y versus MA ($p < 0.01$)* |  |  |  |
| Isoform 2 of Glycerol-3-phosphate dehydrogenase, cytoplasmic (GPD1) | 1984 ± 263 | 1067 ± 510 | 0.0026 |
| Protein NDRG2 (NDRG2) | 1428 ± 200 | 903 ± 268 | 0.0031 |
| Acylphosphatase-2 (ACYP2) | 564 ± 142 | 329 ± 92 | 0.0076 |
| Aspartate aminotransferase, cytoplasmic (GOT1) | 478 ± 54 | 94 ± 48 | 0.0068 |
| Isoform 3 of Exportin-2 (CSE1L) | 443 ± 79 | 256 ± 102 | 0.0031 |
| Beta-enolase (ENO3) | 378 ± 63 | 240 ± 49 | 0.0024 |
| Acyl-coenzyme A thioesterase 2, mitochondrial (ACOT2) | 351 ± 61 | 195 ± 84 | <0.0001 |
| Aspartate aminotransferase, mitochondrial (GOT2) | 267 ± 87 | 57 ± 33 | 0.0001 |
| GTP:AMP phosphotransferase AK3, mitochondrial (AK3) | 264 ± 54 | 132 ± 54 | 0.0004 |
| Adenylosuccinate lyase (ADSL) | 239 ± 22 | 135 ± 52 | 0.0033 |
| Acyl-coenzyme A thioesterase 1 (ACOT1) | 202 ± 61 | 85 ± 42 | 0.0006 |
| Guanidinoacetate N-methyltransferase (GAMT) | 202 ± 50 | 97 ± 22 | 0.0004 |
| Isoform 3 of UV excision repair protein RAD23 homolog A (RAD23A) | 161 ± 37 | 93 ± 28 | 0.0061 |
| Malate dehydrogenase, cytoplasmic (MDH1) | 140 ± 53 | 47 ± 27 | 0.0046 |
| Carboxymethylenebutenolidase homolog (CMBL) | 131 ± 23 | 70 ± 25 | 0.0041 |
| *Top 15 of 447 non-MyoF proteins higher in MA versus Y (p<0.01)* |  |  |  |
| Phospholamban (PLN) | 1212 ± 262 | 2378 ± 572 | 0.0024 |
| Calsequestrin-1 (CASQ1) | 1281 ± 218 | 2344 ± 670 | 0.0082 |
| Trifunctional enzyme subunit beta, mitochondrial (HADHB) | 861 ± 98 | 1561 ± 397 | 0.0041 |
| Trifunctional enzyme subunit alpha, mitochondrial (HADHA) | 756 ± 100 | 1292 ± 335 | 0.0075 |
| Isoform 2 of Sarcalumenin (SRL) | 296 ± 69 | 593 ± 181 | 0.0074 |
| Troponin C (TNNC1) | 291 ± 46 | 494 ± 123 | 0.0071 |
| Troponin I (TNNI1) | 219 ± 28 | 466 ± 145 | 0.0047 |
| Myomesin-2 (MYOM2) | 95 ± 36 | 349 ± 127 | 0.0020 |
| PRA1 family protein 3 (ARL6IP5) | 214 ± 36 | 322 ± 58 | 0.0059 |
| Tropomyosin alpha-3 chain (TPM3) | 129 ± 21 | 295 ± 76 | 0.0011 |
| Complement C1q subcomponent subunit C (C1QC) | 91 ± 8 | 251 ± 49 | <0.0001 |
| Calpain-1 catalytic subunit (CAPN1) | 155 ± 11 | 213 ± 26 | 0.0013 |
| Complement C1q subcomponent subunit B (C1QB) | 74 ± 9 | 206 ± 48 | 0.0002 |
| Mitochondrial import receptor subunit TOM40 homolog (TOMM40) | 83 ± 13 | 206 ± 63 | 0.0023 |
| Isoform 3 of Hexokinase-1 (HK1) | 134 ± 28 | 204 ± 29 | 0.0030 |

Bioinformatics of the non-MyoF proteins that were significantly different between the Y versus MA participants are presented in Table 4 below. Notably, more pathways were predicted to be upregulated in MA versus Y participants.

**Table 4.** Biological processes affected based on non-MyoF proteins different between Y versus MA participants. Legend: Biological pathways presented to be up- and down-regulated between age groups based on differential non-MyoF protein expression differences between age cohorts (i.e., Table 3 data).

| PANTHER GO-Slim Biological process (GO ID) | Proteins altered (pathway proteins) | p-value |
| --- | --- | --- |
| *From 96 non-MyoF proteins higher in Y versus MA, pre- (p<0.01)* |  |  |
| Organophosphate metabolism (GO:0019637) | 14 (395) | <0.0001 |
| *From 447 non-MyoF proteins higher in MA, pre- versus Y (p<0.01)* |  |  |
| mRNA export from nucleus (GO:0006406) | 8 (33) | 0.0042 |
| lipid oxidation (GO:0034440) | 7 (30) | 0.0226 |
| regulation of RNA splicing (GO:0043484) | 10 (75) | 0.0270 |
| protein import (GO:0017038) | 11 (89) | 0.0195 |
| translational elongation (GO:0006414) | 27 (254) | <0.0001 |
| tRNA metabolic process (GO:0006399) | 12 (113) | 0.0317 |
| mitochondrion organization (GO:0007005) | 15 (153) | 0.0071 |
| ubiquitin-dependent protein catabolic process (GO:0006511) | 21 (294) | 0.0102 |
| cellular response to stress (GO:0033554) | 27 (480) | 0.0318 |

### Alternative proteins isoforms in the MyoF and non-MyoF fractions of MA and Y

The enhanced depth of detection provided by proteomics revealed the presence of numerous isoforms in both protein fractions; specifically, there were 175 isoforms for 82 MyoF proteins and 375 isoforms for 173 non-MyoF proteins. The MyoF proteins with the most isoforms included titin (TTN), myosin-binding protein C (MYBPC), and MICOS complex subunit MIC60 (IMMT); each of these targets had four isoforms detected. Given the vast research interest in titin (38, 39), associated isoform data are plotted in Fig. 3a; notably no significant aging or training effects were noted (p>0.01 for all comparisons). The non-MyoF proteins with the most isoforms included Gelsolin (GSN, four isoforms), IMMT (4 isoforms), and Reticulon-4 (4 isoforms); again, no significant age effects were noted for these targets (data not plotted).

**Figure 3.**
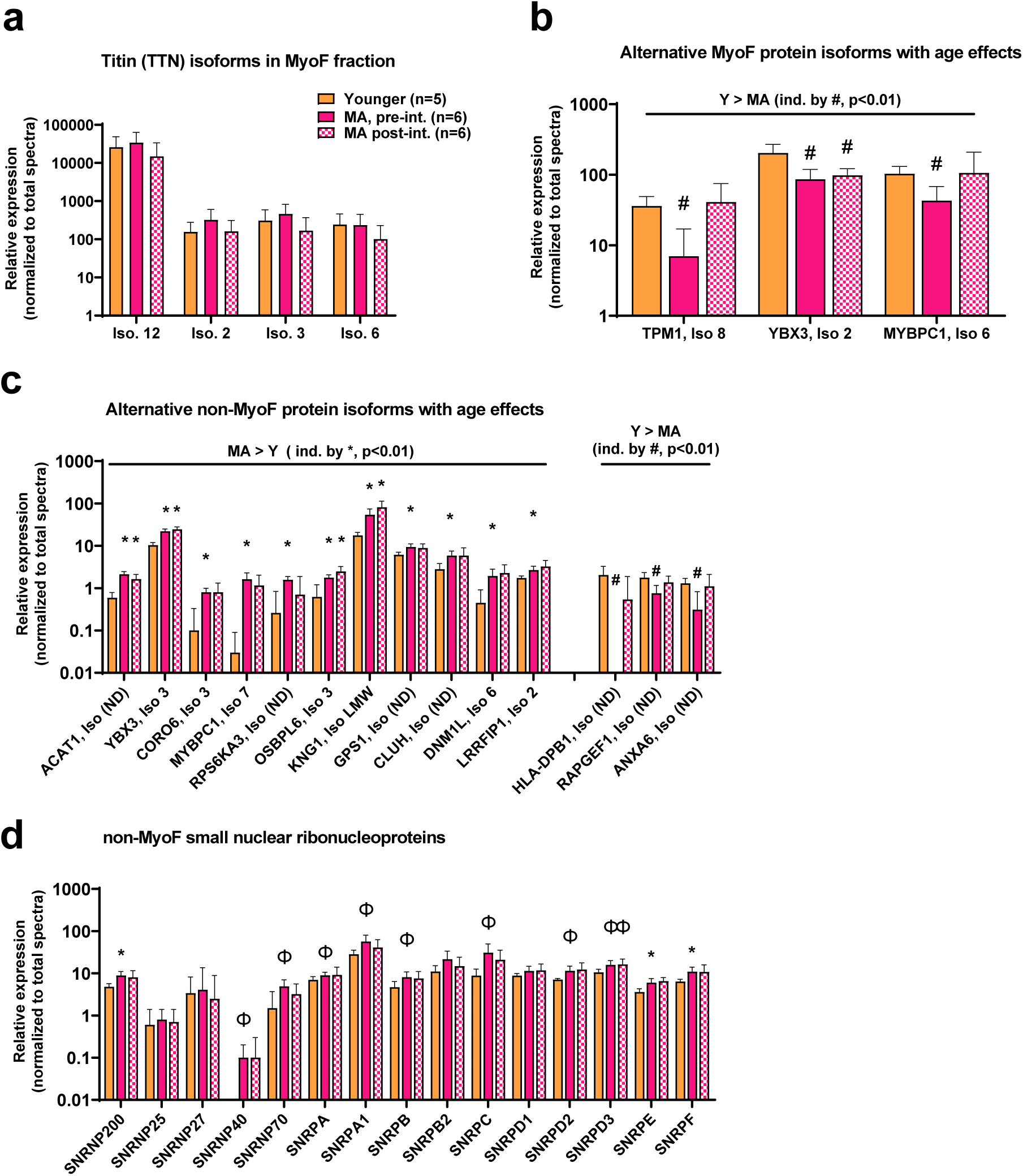
MyoF and non-MyoF alternative protein isoform differences detected with proteomics. Legend: Data presented for Y and MA (pre- and post-intervention) include the identified titin isoforms in the MyoF fraction (panel a), alternative MyoF protein isoforms affected by aging (panel b), and alternative non-MyoF protein isoforms affected by aging (panel c), and small nuclear ribonucleoproteins that make up spliceosomes between cohorts (panel d). Data are presented as mean ± standard deviations for individual protein spectra values (normalized to total run spectra values) and y-axes were scaled as log_10_ for improved visualization. Symbols: #, indicates lower in MA versus Y at one or both time points (p<0.01); *, indicates greater in MA versus Y at one or both time points (p<0.01); Φ, indicates greater in MA versus Y at one or both time points for panel d only (p<0.05). Notes: (ND), indicates that the isoform number was not provided from the Uniprot’s Homo Sapiens reference database (UP000005640_9606).

We plotted significantly different alternative protein isoform abundances between Y and MA in both protein fractions given that bioinformatics on the non-MyoF fraction indicated “regulation of RNA splicing” (GO:0043484) was predicted to be upregulated in the older cohort (Fig. 3b/c). MA (pre-intervention) versus Y comparisons indicated that only three alternative MyoF protein isoforms were different between age groups (all higher in Y, p<0.01). However, 14 alternative non-MyoF protein isoforms were significantly different between age groups (11 higher in MA, p<0.01), and while not depicted in Fig. 3c, 32 additional alternative non-MyoF protein isoforms were numerically different between age groups (25 higher in MA, p<0.05). Further, when examining the abundances of small nuclear ribonucleoproteins belonging to spliceosome complexes in the non-MyoF fraction, three reached the p<0.01 significance threshold as being more enriched in MA (pre-intervention) versus Y participants (SNRP200, SNRPE, SNRPF), and several others were numerically greater in MA participants (SNRP40/70/A/A1/B/C/D2/D3, p<0.05; Fig. 3d). Notably, training did not alter the expression of any SNRP in Fig. 3d (p>0.05 for all), and most of these proteins were not detected in the MyoF fraction.

### MyoF and non-MyoF protein differences prior to and following resistance training in MA participants

In MA, knee extensor resistance training significantly altered 13 MyoF proteins (11 upregulated and two downregulated, p<0.01; Fig. 4a), and 64 non-MyoF proteins (56 upregulated and eight downregulated, p<0.01; Fig. 4b/c). These alterations represented ∼0.3% of the MyoF proteome and ∼1.0% of the non-MyoF proteome. Bioinformatics within each fraction were attempted, albeit no pathways were predicted to be significantly affected.

**Figure 4.**
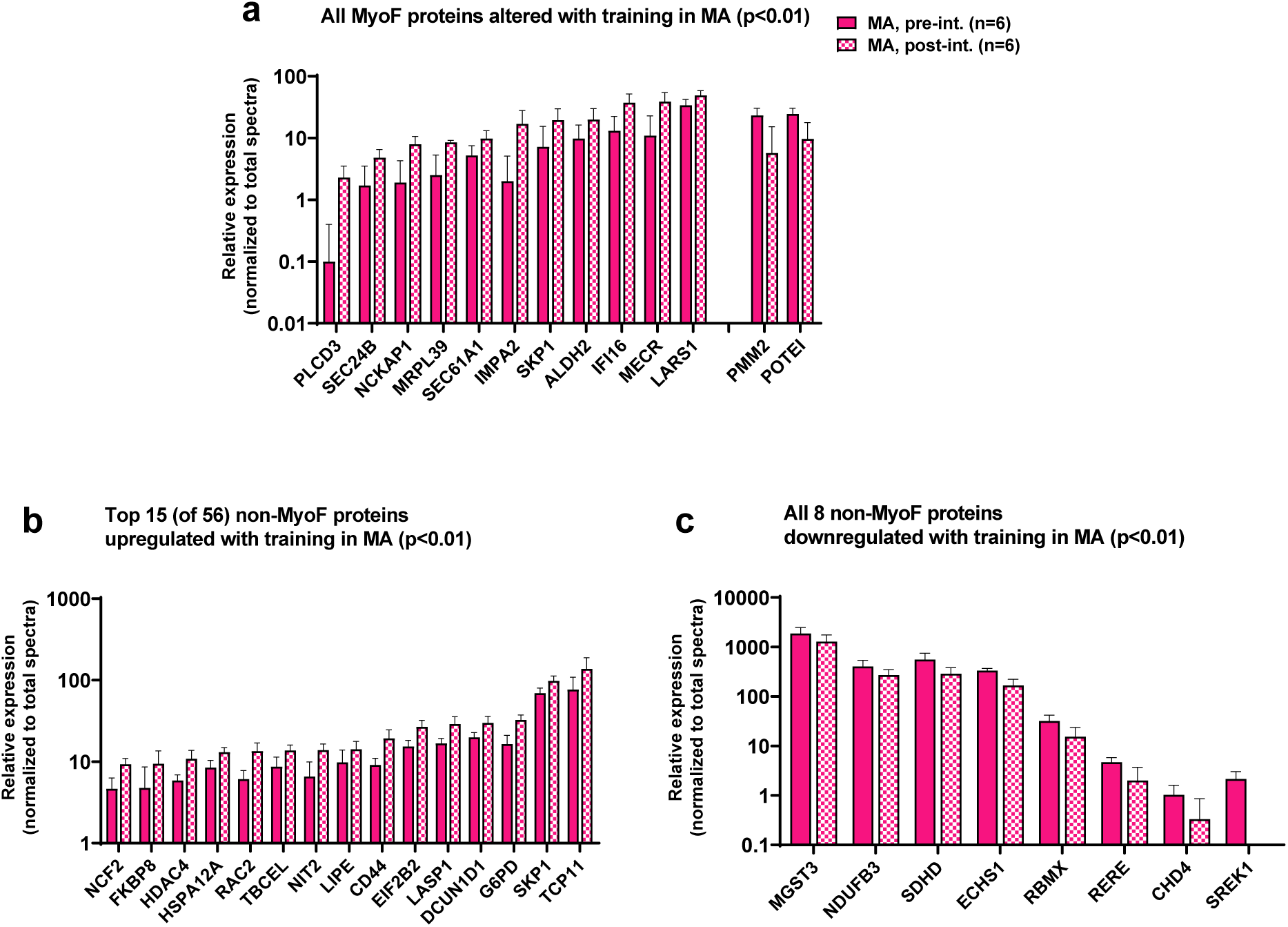
MyoF and non-MyoF proteins altered with resistance training in MA participants. Legend: Data presented for MA prior to and following eight weeks of knee extensor training include proteins in the MyoF fraction (11 up-regulated, 2 down-regulated; panel a), the top 15 up-regulated proteins in the non-MyoF fraction (panel b), and all 8 down-regulated proteins in the non-MyoF fraction (panel c). Data are presented as mean ± standard deviations for individual protein spectra values (normalized to total run spectra values), and y-axes were scaled as log_10_ for improved visualization. Notes: No biological processes were predicted to be affected with training based on these alterations.

### Proteolysis targets manually interrogated in both fractions

Based on bioinformatics indicating that proteostasis was predicted to be altered with aging (Table 3), we manually interrogated proteolysis-related protein targets (i.e., calpain-1/2, and the summed spectra of 26S proteasome subunits). Figure 5 contains these targets including the 26S proteasome subunits detected in the MyoF and non-MyoF fractions (panels a/d), calpain-1 (panels b/e) and calpain-2 (panels c/f). The summed spectra of the 34 detected 26S proteasome subunits in the MyoF fraction was significantly lower in MA at both time points versus Y participants. Both calpains were also numerically lower in the MyoF fraction of MA at both time points versus Y participants (CAPN2 was significant in MA pre-intervention versus Y, p<0.01), and training did not significantly affect either protein. In the non-MyoF fraction, the summed spectra of the 42 detected 26S proteasome subunits was not significantly different between Y versus MA participants at either time point. However, both calpains were higher in the non-MyoF fraction of MA at both time points versus Y participants (CAPN1 was significant in MA pre-intervention versus Y, p<0.01), and training did not significantly affect either protein.

**Figure 5.**
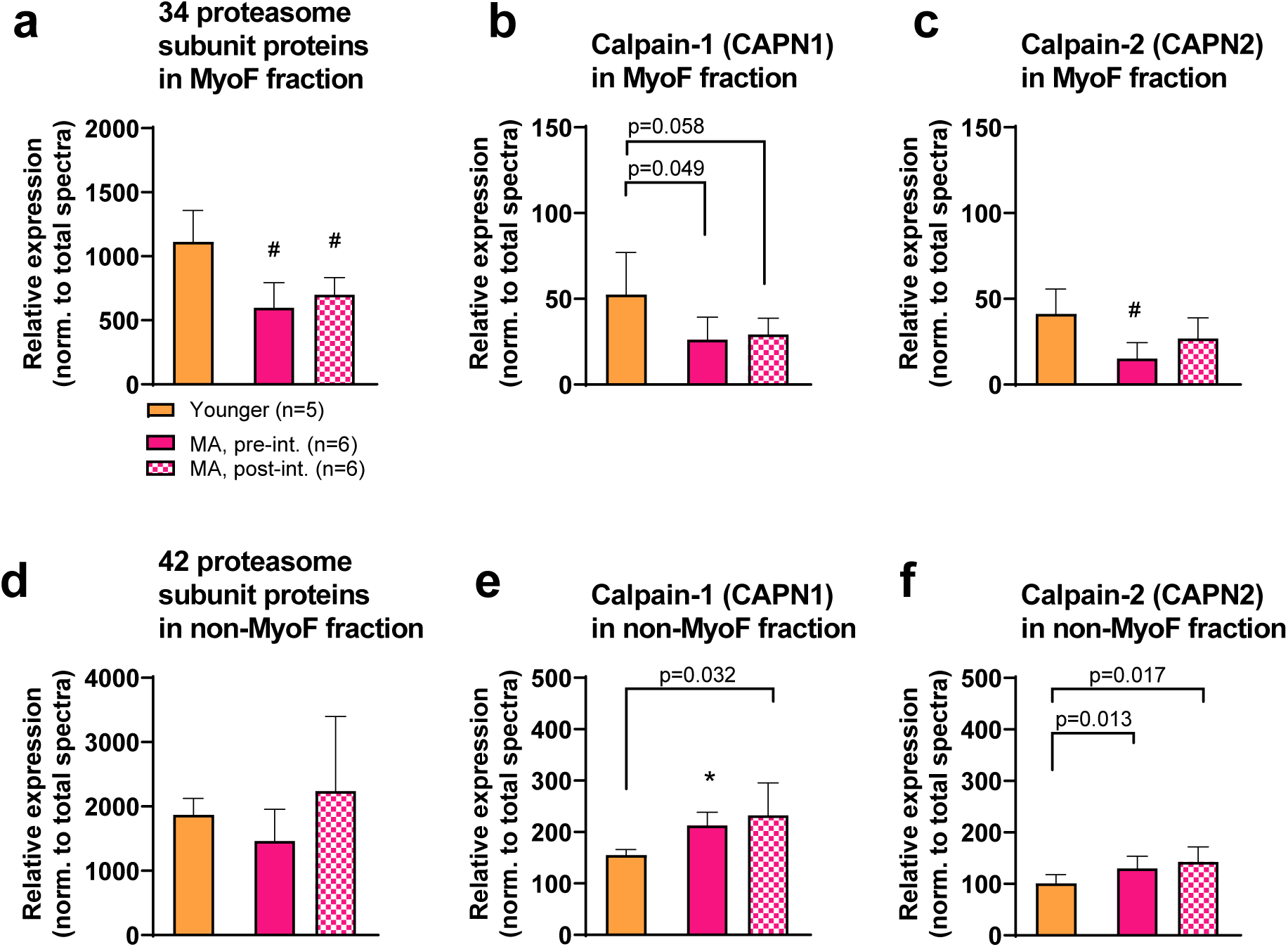
MyoF and non-MyoF proteasome and calpain proteins. Legend: Data presented for Y and MA prior to and following eight weeks of knee extensor training include proteasome subunits and calpains 1/2 in the MyoF fraction (panels a-c) and non-MyoF fraction (panels d-f). Data are presented as mean ± standard deviations for individual protein spectra values (normalized to total run spectra values). Symbols: #, indicates lower in MA versus Y at one or both time points (p<0.01); *, indicates greater in MA versus Y at one or both time points (p<0.01). Notes: Certain p-values not meeting the significance criteria for these data were presented due to the visual differences observed between cohorts.

## DISCUSSION

Using a novel analytical approach, we examined the deep proteomic signatures of the MyoF and non-MyoF fractions in younger adults as well as middle-aged participants before and after eight weeks of knee extensor resistance training. More non-MyoF proteins differed between age cohorts compared to MyoF proteins (8.4% versus 2.5% of the respective protein pools). More non-MyoF proteins (447/543) were also more highly abundant in MA versus Y and bioinformatics predicted that several biological processes were more operative in the older participants. A greater abundance in alternative variants, proteins associated with spliceosomes, and proteolysis-related proteins were also evident in the non-MyoF fraction of MA versus Y, and these observations corroborated certain bioinformatics findings. Although resistance training in MA non-significantly increased VL cross-sectional area (+6.5%) and significantly increased knee extensor strength (+8.7%), training marginally affected the MyoF and non-MyoF proteomes and no biological processes were predicted to be affected in either fraction. These findings will be expanded upon in the paragraphs below.

As stated above, several studies have performed proteomic analyses on skeletal muscle to compare molecular signatures that exist between younger and older adults or to examine how resistance training affects this aspect of the muscle-molecular milieu (1, 14, 18–20, 22). The novelty of the current study was the proteomics approach utilized and the knowledge gained relative to these prior investigations. Fractionation of muscle into solubilized MyoF and non-MyoF homogenates enabled the detection of unique proteins in each fraction, which has only been attempted in one other study to our knowledge (1). In this prior study, we performed bottom-up LC-MS-based proteomics on each fraction from younger resistance-trained, younger untrained, and older untrained men (n=6 per group). We identified a total of 810 proteins in both fractions that were expressed in at least one participant. In the current study we detected a total of 10,866 proteins in both fractions. While most of identified proteins were present in both fractions, we were able to identify 2,217 unique non-MyoF proteins and 193 unique MyoF proteins. This robust increase in detection depth (∼13.4-fold) is insightful for numerous reasons. First, it was revealed that metabolic enzymes constituted the top class of proteins in both fractions as well as proteins that overlapped in both fractions. Hence, although myosin heavy chain isoforms, troponin, titin, and actin were the most enriched in the MyoF fraction, these data counter the notion that the MyoF fraction contains mainly contractile proteins. Several nuclear proteins were also identified in the MyoF fraction (e.g., histones and other chromatin-binding proteins) indicating that our MyoF isolation method likely pellets nuclei. Finally, we were able to identify numerous proteins that are not commonly reported in previous skeletal muscle proteomic studies (see Table 1 and Fig. 2 for example). To this end, several MYH isoforms beyond the three common 7/2/1 isoforms were highly enriched in the MyoF fraction, and this may be due to the persistence of non-conventional or developmental isoforms in certain regions of adult myofibers or in transitioning fibers as discussed by Schiaffino et al. (40). Both fractions contained most of the large (∼40) and small (∼30) ribosomal subunit proteins, mitochondrial oxidative phosphorylation enzymes, and mitochondrial ribosomal proteins (>70), all of which likely represents the presence of sarcoplasmic and intermyofibrillar mitochondria. Both fractions also contained numerous alternative isoforms for several proteins, various transcription factors (e.g., MEF2D, F-box proteins, SMAD1/2/3, NFAT isoforms, and several others), DNA and RNA polymerase subunits, various growth factors and their receptors (e.g., EGFR, VEGFA, FGF2/13, PDGFRa/b, TGFB1/2, and others), dozens of eukaryotic initiation/elongation factors, nearly 50 solute carrier family member proteins (i.e., nutrient and metabolite transporters), cytoplasmic and mitochondrial aminoacyl tRNA ligases, and hundreds of signal transduction proteins of interest to skeletal muscle biologists (e.g., mTOR, RPTOR, p70s6k, AMPK subunits, cyclin-dependent kinases and inhibitors, and others). We believe this enhanced level of detection was due to muscle fractionation, and more importantly, the utilization of NPs prior to mass spectrometry. Indeed, this same contention has been posited by others using this technology to increase detection depth of circulating proteins in human plasma (25, 41).

Notable MyoF and non-MyoF proteome signatures between age groups were also evident. For instance, aging seemingly affects the non-MyoF protein signature more so than the MyoF fraction. This finding agrees with our past proteomic study where we reported that 37 non-MyoF proteins (versus only 18 MyoF proteins) were differentially expressed between college-aged and older men (mean age 62 years old) (1). However, the increased detection depth in the current study indicated that 112 of the 4,421 identified MyoF proteins met the p<0.01 significance threshold between age cohorts. Moreover, all but one of these proteins (TMPO, Lamina-associated polypeptide 2 isoforms beta/gamma) were greater in Y versus MA indicating either a loss or decreased expression of ∼2.5% proteins belonging to the MyoF fraction. Although no biological processes were predicted to be affected between age cohorts based on this list of 111 proteins, several of these targets were notable. For example, three heat shock proteins (HSPB1, HSP90AB1, HSPD1) were more lowly abundant in MA participants and this agrees in principle with past literature indicating the expression of heat shock proteins in skeletal muscle is dysregulated with aging (42, 43). Several MyoF mitochondrial proteins were also lower in MA participants (DECR1, NDUFS8, ETFDH, GPD2, STOML2, ALDH2, COQ5, IARS2, ALDH1B1, MRPL32). This also agrees with past literature indicating either a decrease in mitochondrial volume density or decreased mitochondrial function with skeletal muscle aging (44), and more specifically agrees with a study by Callahan et al. (45) who reported that older participants presented reductions in the size of intermyofibrillar mitochondria.

Strikingly, 543 proteins (∼8.4%) of the non-MyoF proteome were different between MA and Y participants, and unlike the trends observed in the MyoF fraction, most of these proteins (447) were significantly enriched in the MA cohort. These figures agree with a proteomic investigation by Robinson et al. (14) who reported that more muscle proteins (220/347) were higher in untrained older versus younger individuals prior to a period of exercise training. These robust differences between age cohorts in the current study also revealed that several biological processes were predicted to be upregulated in MA participants. Some of these processes either contradict each other or agree with past literature reporting similar aging phenotypes. Regarding the former, while more proteins associated with translation elongation were more abundant in MA participants (which would potentially promote muscle anabolism), proteins associated with ubiquitin-mediated proteolysis were also more enriched. One interpretation of these data could be that aging increases skeletal muscle protein turnover. However, this is likely not the case given that a variety of studies ranging from human tracer studies to nematode models have indicated that protein turnover in response to feeding or in a basal state is impaired with aging (46, 47). This aspect of our data also agrees with a report by Ubaida-Mohien et al. (22) who showed that 31% of proteins related to proteostasis were altered with age (24 underrepresented and 50 overrepresented, p<0.05) in healthy older versus younger adults. Hence, we posit that proteins associated with these processes may have been more abundant in MA participants in a compensatory attempt to counter age-related declines in muscle protein turnover. It is also interesting that proteasome subunits and calpains were more enriched in the MyoF fraction of Y versus MA, whereas these same proteins were more enriched in the non-MyoF MA versus Y. While speculative, an enrichment of proteolytic proteins in the MyoF fraction might play a role in functional proteostasis, while an enrichment of these proteins in the non-MyoF fraction might be indicative of a gradual dysregulation in proteostasis.

Non-MyoF proteins associated with mRNA export and splicing were also elevated in MA participants, and this agrees with other reports. For instance, a recent review by Park et al. (48) cites a variety of cell culture evidence to suggest that the nuclear pore complex is disrupted with aging and that this leads to a dysregulation in mRNA export. A rodent study by Mobley et al. (49) also suggests that mRNA levels linearly decrease in skeletal muscle with increasing age. Hence, again, a higher abundance in non-MyoF proteins associated with mRNA export could also be a compensatory response to offset these age-associated effects burdened by myonuclei. The greater abundance of proteins associated with mRNA splicing is striking and agrees in principle with a report by Rodriguez et al. (50) who showed that the skeletal muscle of aged mice possessed ∼4 times more RNA splice variants than younger counterparts. Our data also agree with the abovementioned proteomics report by Ubaida-Mohien et al. (22) who showed that proteins related to alternative splicing were more abundant in healthy older (versus younger) adults. We also performed a follow-up analysis showing that several small nuclear ribonucleoproteins (SNRPs, or snRNPs) that make up spliceosomes and alternative non-MyoF protein isoforms (indicative of increased spliceosome activity) were elevated in MA versus Y participants. This is particularly insightful given that dysfunctional spliceosome activity and the aberrant RNA and protein expression of splice variants have been linked to age-associated maladies such as cellular senescence (51, 52). Hence, these independent reports of age-associated increases in splice RNA and protein variants, along with the current data suggesting that the relative abundances of spliceosomes are greater in older participants, warrant future research elucidating the causes and consequences of this phenomenon.

A final noteworthy topic was the observation of marginal MyoF and non-MyoF alterations with eight weeks of knee extensor training in MA participants. Although this may have been due to the modest training regimen that only lasted eight weeks in duration, limited proteome plasticity with aging and/or the limited ability of resistance training to alter the muscle proteome could also be plausible explanations. Support for both phenomena come from Robinson et al. (14) who employed deep proteomics to report that ∼200 muscle proteins were altered in older participants after 12 weeks of resistance training (p<0.05), and this was less than the ∼300 proteins that were altered in the younger participants. Hence, an aging effect was noted. However, assuming the authors identified >3,000 muscle proteins, which was not reported to our knowledge, this represents less than 10% of the detectable proteome being altered with resistance training. Deane et al. (13) used a different proteomics approach to examine the non-MyoF proteome adaptations in older participants following 20 weeks of resistance training in younger and older adults. Although their depth of detection was limited to ∼160 proteins, resistance training only increased five non-MyoF proteins in older participants when a p<0.05 significance threshold was employed (i.e., ∼3% of the detectable proteome). Interestingly, this effect was not confined to older participants given that the younger participants in their study only presented an elevation in four non-MyoF proteins with training. Our laboratory also used proteomics to examine non-MyoF protein adaptations in college-aged men following ten weeks of resistance training (1). Only 13 proteins were shown to be altered with training (12 up, one down, p<0.05) and this represented ∼3.4% of the detectable non-MyoF proteome. Hence, these two latter studies do not support the aging hypothesis and, instead, provide evidence of limited muscle proteome plasticity with resistance training. Despite marginal non-MyoF proteome alterations in MA participants with training, there were interesting targets that were altered. For instance, the UBR7 E3 ligase was upregulated, and recent evidence suggests that an E3 ligase in this same protein family (UBR5) is required for load-induced skeletal muscle hypertrophy (53). Additionally, the knockdown of another member of this family (UBR4) promotes hypertrophy in *Drosophila* and mice (54). SRC was upregulated and this non-receptor tyrosine kinase has been implicated in interacting with vitamin D to promote anabolic signaling in skeletal muscle (55). HDAC4 was upregulated and this adaptation could be operative in muscle-metabolic adaptations and ultrastructural remodeling to resistance training given that HDACs have been implicated in controlling the expression of various metabolic and contractile protein genes (56, 57). Two protein phosphatases were also upregulated (PPP1R12A and PIP4P2), and both have been shown to be involved in aspects of insulin and growth factor signaling (58, 59).

Indeed, this study is limited by the limited MA and Y sample sizes, and Y participants being women. Furthermore, the lack of training data in Y participants to examine age-associated effects is a limitation, and we lacked remaining skeletal muscle to perform downstream analyses (e.g., examining RNA splice variants and/or proteasome activity assays) which may have provided additional insight. However, a primary objective of this publication was to feature our novel proteomic approach as we believe that this will add tremendous insight into the field of skeletal muscle biology. Likewise, the data from this study can be used to generate hypotheses for other age-related or resistance training proteomic or targeted protein approaches moving forward.

In conclusion, we provide preliminary evidence to support that muscle aging predominantly affects the non-MyoF protein pool and that this is associated with biological processes which may act to counteract dysfunctional cellular homeostasis. We also provide preliminary evidence of limited MyoF and non-MyoF proteome plasticity to shorter-term resistance training in middle-aged participants, and this agrees with prior proteomic investigations. Finally, and most importantly, we believe that the utilization of skeletal muscle tissue fractionation protocols and NP-based protein corona formation prior to downstream proteomics has the potential to add incredible insight in identifying novel protein targets affected by exercise training, aging, and various disease states.

## ACKNOWLEDGEMENTS

We thank the participants who volunteered and participated in the study. We also thank Seer, Inc. and their team members (David Hill, Mara Riley, Ryan Hill, Aaron S. Gajadhar, Khatereh Motamedchaboki, and others) for assisting us through the various procedures needed to perform proteomics analysis including performing pilot feasibility experiments using the Proteograph Product Suite. Shao-Yung Chen is an employee of Seer, Inc. who was largely responsible for the execution of the pilot feasibility experiments, although he had no role in the study design. None of the other co-authors have apparent conflicts of interest in relation to these data.

## DATA AVAILABILITY

Raw data related to the current study outcomes will be provided upon reasonable request by emailing the corresponding author.

